# Introgression underlies phylogenetic uncertainty but not parallel plumage evolution in a recent songbird radiation

**DOI:** 10.1101/2023.04.25.538255

**Authors:** Loïs Rancilhac, Erik D. Enbody, Rebecca Harris, Takema Saitoh, Martin Irestedt, Yang Liu, Fumin Lei, Leif Andersson, Per Alström

## Abstract

Instances of parallel phenotypic evolution offer great opportunities to understand the evolutionary processes underlying phenotypic changes. However, confirming parallel phenotypic evolution and studying its causes requires a robust phylogenetic framework. One such example is the “black-and-white wagtails”, a group of five species in the songbird genus *Motacilla*: one species, the White Wagtail (*M. alba*), shows wide intra-specific plumage variation, while the four others form two pairs of very similar-looking species (African Pied Wagtail *M. aguimp* + Mekong Wagtail *M. samveasnae* and Japanese Wagtail *M. grandis* + White-browed Wagtail *M. maderaspatensis*, respectively). However, the two species in each of these pairs were not recovered as sisters in previous phylogenetic inferences. Their relationships varied depending on the markers used, suggesting that gene tree heterogeneity might have hampered accurate phylogenetic inference. Here, we use whole genome resequencing data to explore the phylogenetic relationships within this group, with a special emphasis on characterizing the extent of gene tree heterogeneity and its underlying causes. We first used multispecies coalescent methods to generate a “complete evidence” phylogenetic hypothesis based on genome-wide variants, while accounting for incomplete lineage sorting and introgression. We then investigated the variation in phylogenetic signal across the genome, to quantify the extent of discordance across genomic regions, and test its underlying causes. We found that wagtail genomes are mosaics of regions supporting variable genealogies, because of ILS and inter-specific introgression. The most common topology across the genome, supporting *M. alba* and *M. aguimp* as sister species, appears to be influenced by ancient introgression. Additionally, we inferred another ancient introgression event, between *M. alba* and *M. grandis*. By combining results from multiple analyses, we propose a phylogenetic network for the black-and-white wagtails that confirms that similar phenotypes evolved in non-sister lineages, supporting parallel plumage evolution. Furthermore, the inferred reticulations do not connect species with similar plumage coloration, suggesting that introgression does not underlie parallel plumage evolution in this group. Our results demonstrate the importance of investigation of genome-wide patterns of gene tree heterogeneity to help understanding the mechanisms underlying phenotypic evolution.

## 1 Introduction

Phylogenetic studies of evolutionary radiations provide the foundation for studying the processes underlying speciation and species diversification. Accurately inferring evolutionary relationships among lineages is needed to characterize patterns of phenotypic evolution, such as parallel evolution (Elmer and Meyer 2011; Stern 2013), and to understand large-scale evolutionary processes. Factors that complicate phylogenetic inference, including standing variation and introgression, are increasingly appreciated for their role in generating novel phenotypes and contributing to speciation (Baack and Rieseberg 2007; Enciso-Romero et al. 2017; Stryjewski and Sorenson 2017; Marques et al. 2019; Svardal et al. 2020). Studying diversification from this point of view requires a well resolved phylogenetic framework and an understanding of the relative contribution of these factors to the species genetic diversity.

Species tree inference in rapid radiations is complicated because the prevalence of Incomplete Lineage Sorting (ILS) increases as branches get shorter (Nichols 2001; Degnan and Rosenberg 2009), and closely related species often experience gene flow throughout the divergence process (and sometimes post-divergence (Seehausen 2004; Rheindt and Edwards 2011; Lamichhaney et al. 2015). More generally, introgression can contribute to novel phenotypes in rapid adaptive radiations (Dasmahapatra et al. 2012; Wallbank et al. 2016; Enciso-Romero et al. 2017; Han et al. 2017; Marques et al. 2019; Svardal et al. 2020). For these reasons, rapid radiations are very likely to feature high levels of gene-tree heterogeneity (Giarla and Esselstyn 2015; Cai et al. 2021; Astudillo-Clavijo et al. 2022). ILS has long been recognized as a confounding factor in phylogenetic inference, and it is now standard to take it into account by analyzing multi-locus data through Multispecies Coalescent (MSC) models (Rannala and Yang 2003; Maddison and Knowles 2006). Distinguishing the respective contributions of ILS and introgression is more challenging, but this issue has gathered widespread attention in the past decade, and a variety of methods have been developed to that end (reviewed by Hibbins & Hahn, 2022a). Yet, characterizing introgression events remains challenging in the presence of phylogenetic uncertainty (e.g., when ILS introduces background noise; Beckman et al., 2018; Pease, 2018), especially when using tests based on excess of allele sharing between non-sister species, whose interpretation rely on assumptions on the species tree (Durand et al. 2011). This results in a paradox for evolutionary biologists: introgression contributes to phylogenetic uncertainty, but identifying introgressed gene trees becomes more difficult as phylogenetic uncertainty increases (Hibbins and Hahn 2022b).

Beyond the field of phylogenetics, separating the true species tree from the alternative topologies generated by ILS and introgression has two important consequences for evolutionary biology research. First, identifying species relationships is a prerequisite to understand whether similar phenotypes evolved from the same ancestor or independently (Elmer and Meyer 2011). Second, when similar phenotypes are shared across non-sister branches of the species tree, understanding the drivers of gene tree heterogeneity is useful to determine whether the variants underlying focal phenotypes represent standing variation (ILS), were transferred horizontally between branches (introgression), or are lineage-specific *de novo* variants (Pease et al. 2016; Marques et al. 2019; Svardal et al. 2020; Alaei Kakhki et al. 2023).

Here, we explore this issue using the “black-and-white wagtails”, a monophyletic group of recently diverged species (ca 0.7–2 million years ago [Ma]; Alström et al., 2015; Harris et al., 2018) in the avian genus *Motacilla* (Fig. 1). This group includes the White (*M. alba*), African Pied (*M. aguimp*), Japanese (*M. grandis*), White-browed (*M. maderaspatensis*) and Mekong (*M. samveasnae*) wagtails (Alström and Mild 2003). *Motacilla alba* shows pronounced variation across its range, with nine subspecies (Fig. 1b), whereas the four other species show little or no intra-specific geographical variation but form two species pairs with very similar plumages: *M. aguimp* and *M. samveasnae* are rather small species with a white throat and white patch on the side of the neck; *M. grandis* and *M. maderaspatensis* are larger, with a black throat and neck side (Fig. 1). Prior analyses of reduced-representation genome-wide data recovered the black-and-white wagtails as monophyletic. However, although the relationships among species changed among analyses, none of the proposed phylogenetic hypotheses recovered phenotypically similar species as sisters (Harris et al. 2018). Phylogenetic conflicts between mitochondrial and nuclear markers, as well as between independent nuclear loci, suggest that these relationships may not represent the “true” species tree (Alström & Ödeen 2002; Drovetski et al. 2018; Harris et al. 2018). Two alternative hypotheses could explain this pattern: 1) species with similar phenotypes are sister in the species tree, but previous inferences were obscured by gene tree heterogeneity; or 2) similar phenotypes evolved in non-sister branches of the tree, possibly facilitated by retention of ancestral polymorphism or introgression.

**Figure 1.**
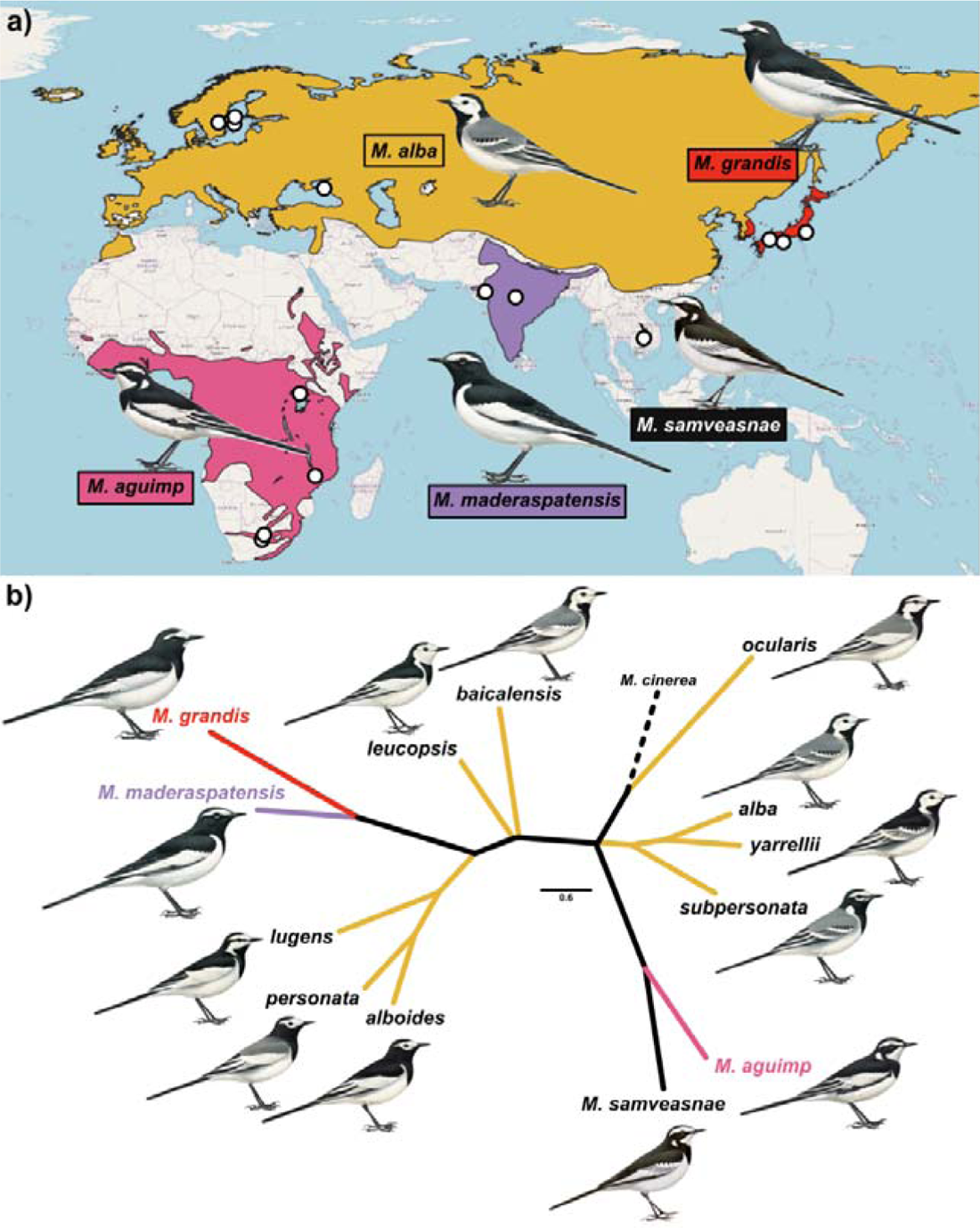
Sampling and plumage variation. a) Distribution of the five black-and-white wagtail species (BirdLife International and Handbook of the Birds of the World 2021), with sampling localities shown as white circles. b) Unrooted Neighbor-joining tree constructed from 16 plumage characters scored in adult breeding males of all five species, including the nine subspecies of M. alba, plus M. cinerea as an outgroup. The branch leading to M. cinerea has been artificially shortened for improved graphical resolution. Paintings courtesy Bill Zetterström (from Alström and Mild 2003).

We here use whole genome data to evaluate these alternative explanations for phylogenetic discordance by characterizing genome-wide patterns of gene tree heterogeneity, and determine their underlying causes. By interpreting plumage similarity in the light of genome-wide patterns of phylogenetic variation and their underlying causes, we provide a solid basis to understand phenotypic parallelism in the *Motacilla* radiation.

## 2 Material and Methods

### 2.1 Analyses of plumage characteristics

As a basis to investigate parallelism in plumage evolution, we first used plumage characters to quantify the similarity among the five black-and-white wagtail species (including all subspecies of *M. alba*). We scored 16 discrete plumage characters in breeding plumage adult males based on descriptions in Alström and Mild (2003), corresponding to all characters showing little intra-specific variation and segregating at least two taxa (Supplementary material SM1, Table S1). The nine subspecies of *M. alba* were included as a way to verify whether they resemble each other more closely than the four other species. Pairwise distances were calculated on a binary basis, i.e., giving a distance of 0 if the two species have the same character state and 1 if they differ. The obtained matrix was used to calculate a Neighbor-Joining (NJ) tree using the R package phyclust (Chen 2011), with the distantly related Grey Wagtail (*M. cinerea*) included as an outgroup.

### 2.2 Sampling, genomes sequencing, assembly and variant calling

We resequenced genomes for 29 samples covering the five black-and-white wagtail species (*M. aguimp*: N=6, *M. alba*: N=7, *M. grandis:* N=6, *M. maderaspatensis*: N=2, *M. samveasnae*: N=2; Fig. 1A, Table S2) and the outgroup species Grey Wagtail (*M. cinerea*, N=6). Only the nominate subspecies of *M. alba*, which is allopatric with the other species, was sampled to avoid potential phylogenetic noise introduced by recent gene flow with sympatric species (hybridization between *M. alba lugens* and *M. grandis* has been documented in Japan; Alström and Mild 2003). DNA was extracted using the DNeasy Blood and Tissue Kit (Qiagen, Hilden, Germany). Whole genomes were resequenced on an Illumina NovaSeq 6000 S4 (Illumina, CA) at SciLifeLab, Stockholm and Uppsala (Sweden). We removed Illumina adapters from raw reads using fastp (Chen et al. 2018). Trimmed short-reads were aligned to the chromosome level assembly for *Motacilla alba* (GenBank accession GCA_015832195.1) with bwa mem v.0.7.17-r1188 using the accelerated Sentieon engine (Freed et al. 2017). We removed PCR duplicates using the LocusCollector and ‘dup’ engine in Sentieon. We called variable sites in each individual using HaplotypeCaller (GATK v4.1, McKenna et al. 2010), as implemented in the Sentieon tools ‘HaploTyper’ function and setting a genotype quality confidence threshold of 30 (--emit_conf 30). We included high confidence invariant sites in the output to distinguish high confidence homozygous reference sites from uncertain variant calls that were filtered out. We performed joint-genotyping using the GVCFtyper engine in Sentieon tools, which corresponds to the JointGenotyper tool of GATK v4.1. We removed variants that did not pass the following standard GATK filters using the VariantFiltration command: ReadPosRankSum < −0.8, QD < 2, FS > 60.0, SOR > 3.0, MQ < 40.0, and MQRankSum < −12.5. Genotypes with less than 2x sequencing depth, greater than 100x sequencing depth, or less than genotype quality of 20 were set to missing. We next retained only SNPs using GATK SelectVariants. Separately, we retained invariant sites if > 50% sites were determined to be invariant and no SNPs were called. A combined call set was generated by merging both variant and invariant sites.

To infer haplotypes, we used the statistical haplotype phasing software SHAPEIT4 (Delaneau et al. 2019). Prior to analysis we identified phase informative reads for each individual using WhatsHap (Martin et al. 2016). Individual VCFs with phase informative reads annotated were passed to SHAPEIT4 using the –sequencing flag and setting a phase sets threshold (i.e., phase informative read scoring) of 0.0001. For chromosome Z phasing, we only used male samples to avoid the possibility of mis-called heterozygotes in females, the heterogametic sex in birds. We removed missing genotypes that were imputed by SHAPIT4 using bcftools v1.12, so that our variant data set only included variant with read-supported data.

### 2.3 Multispecies coalescent species trees and networks analyses

To simultaneously account for ILS and estimate phylogenetic relationships among the black-and-white wagtails, we used the SNPs-based full-coalescent method implemented in SNAPPER (Stoltz et al., 2021), based on three SNP subsets. First, we selected all bi-allelic SNPs that had no missing genotypes and were phylogenetically informative (minor allele count ≥2) in all 29 samples, thinned them to 5,000 bp to mitigate the effects of linkage disequilibrium (*snapper_5sp* dataset; 167,186 SNPs), and converted them into a binary nexus file using vcf2phylip.py v2.8 (Ortiz 2019). We used this matrix to infer a species tree in SNAPPER v1.0 with default priors. We set up the SNAPPER analyses with an MCMC of 800,000 steps, sampling every 1,000, and assessed convergence with Tracer v1.7 (Rambaut et al. 2018). Because preliminary investigations showed that convergence could not be reach in a reasonable run time, we ran three independent SNAPPER analyses, with different seeds, and combined the results in LogCombiner v2.6.3. The posterior sampling of trees was visualized in Densitree v2.2.7 (Bouckaert 2010). Secondly, because the two samples of *M. maderaspatensis* have only fragmented data (24% and 68% of missing data respectively), we excluded them and reapplied the same filters as above (*snapper_4sp* dataset; 182,693 SNPs). Third, we selected variants located on the Z chromosome for a subset of 18 male samples (Table S2) excluding *M. maderaspatensis*, with the same filtering options as above (14,904 SNPs). For the two *4sp* datasets, a single SNAPPER run with an MCMC of 2,000,000 steps, sampling every 1,000, was enough to reach convergence.

We also performed quartet-based species tree inference using ASTRAL (Mirarab et al. 2014). We inferred phylogenetic trees in non-overlapping 50 kb sliding windows along all autosomes with RAxML v8.2.12 (Stamatakis 2014) using the script raxml_sliding_windows.py from the genomics general package (https://github.com/simonhmartin/genomics_general). For this analysis we included only sites at which *M. maderaspatensis* had less than 50% missing genotypes. We selected windows that 1) were non-adjacent, to reduce the effect of linkage, and 2) had ≥100 SNPs, an arbitrary threshold set to mitigate the effect of low informativeness (*ASTRAL_5sp* dataset; 9,490 trees). As above, gene trees were also inferred by excluding *M. maderaspatensis* and applying the same filters (*ASTRAL_4sp* dataset; 9,443 trees). Both sets of trees were subsequently used as input in ASTRAL-III v5.7.8 (Zhang et al. 2018) for species tree inference.

In order to infer evolutionary relationships in the black-and-white wagtails while accounting for both ILS and introgression, we inferred phylogenetic networks using the coalescent method implemented in SNaQ! (Solís-Lemus and Ané 2016; Solís-Lemus et al. 2017). This method uses the expected distribution of quartet Concordance Factors (CFs) under the multispecies network coalescent to calculate the pseudolikelihood of a phylogenetic network given a set of gene trees or SNPs. We converted the *snapper_4sp* and *snapper_5sp* datasets to phylip format using vcf2phylip v2.8, with heterozygous genotypes randomly resolved. We estimated CFs for all SNPs using SNPs2CF v1.5 (Olave and Meyer 2020), which were used as input for SnaQ!. Phylogenetic network inferences were run with a maximum number of reticulations (*h_max_*) ranging from 0 to 3 and 10 independent searches for each. The best fitting *h_max_* was determined as the highest value generating a significant decrease of the loglik, as recommended by the authors. Out of the ten networks estimated with the selected *h_max_* value, we considered the one with the lowest loglik score to be best fitting our data.

### 2.4 Exploring phylogenetic conflicts

To determine the extent of topological heterogeneity along the genome, we conducted topology weighting analyses in non-overlapping 50 kb sliding windows. For this analysis, we considered all autosomal SNPs but excluded *M. maderaspatensis* to avoid phylogenetic inferences errors due to fragmentary sequences (Sayyari et al. 2017). Phylogenetic trees were calculated in RAxML v8.2.12 (Stamatakis 2014) using the script raxml_sliding_windows.py from the genomics general package. Windows with < 100 SNPs were not considered. Topology weighting was subsequently run in TWISST (Martin and Van Belleghem 2017). As for the species tree analysis, topology weighting was also run separately on the Z chromosome.

Furthermore, we quantified the extent to which our data is affected by ILS and introgression by calculating Patterson’s *D* statistics (Durand et al. 2011; Green et al. 2010) from all autosomal SNPs, in the following four taxon combinations: ((P1=*samveasnae*, P2=*grandis*), P3=*aguimp*); ((P1=*grandis,* P2=*alba*), P3=*aguimp*); ((P1=*samveasnae*, P2=*alba*), P3=*aguimp*); ((P1=*samveasnae*, P2=*grandis*), P3=*alba*). *M. cinerea* was used as outgroup (P4). We also calculated *f_b_* statistics based on two alternative topologies: (*M. aguimp*, (*M. alba*, (*M. samveasnae*, *M. grandis*))) and ((*M. aguimp, M. alba*), (*M. samveasnae, M. grandis*)) (cf. Results). *D* and *f_b_* were calculated in Dsuite v0.5 r45 (Malinsky et al. 2021).

### 2.5 Testing the phylogenetic positions of M. aguimp and M. alba

As our phylogenetic analyses yielded conflicting topologies, we conducted targeted analyses to identify the source of variation among phylogenetic hypotheses. Specifically, the inferred networks induced two placements for *M. aguimp*: in the first (“tree1”), *M. aguimp* was sister to the other four species, while in the second (“tree2”) it was sister to *M. alba* (Fig. 2c, Fig. S4; see Results for more details). To determine which of these trees best represent the species relationships (i.e., the history of speciation events, as opposed to introgression events), we used a “minimum node height” criterion (Fontaine et al. 2015; Li et al. 2019; Forsythe et al. 2020), based on the following expectation, under the assumption that ancient introgression has affected the topology: as the introgression event along the *M. alba* branch happened after it split from its sister species, the most recent split (T1) in gene trees representing the species tree should be deeper than in those inherited through introgression (Fig. 3d). To test that, we used the *4sp* 50 kb sliding windows trees previously inferred, pruned them to remove *samveasnae,* and categorized them depending on whether they support “tree1” or “tree2”, using a custom R script based on ape v5.6-2 (Paradis & Schliep 2019). We then calculated pairwise nucleotide divergence (Dxy) as a proxy for divergence time between species, using the popgenWindows.py script from the genomics general package. Finally, we compared the distribution of Dxy between *grandis* and *alba* in windows supporting “tree1” (T1.tree1) to the distribution of Dxy between *aguimp* and *alba* in windows supporting “tree2” (T1.tree2) using a non-parametric Wilcoxon test calculated in R.

**Figure 2.**
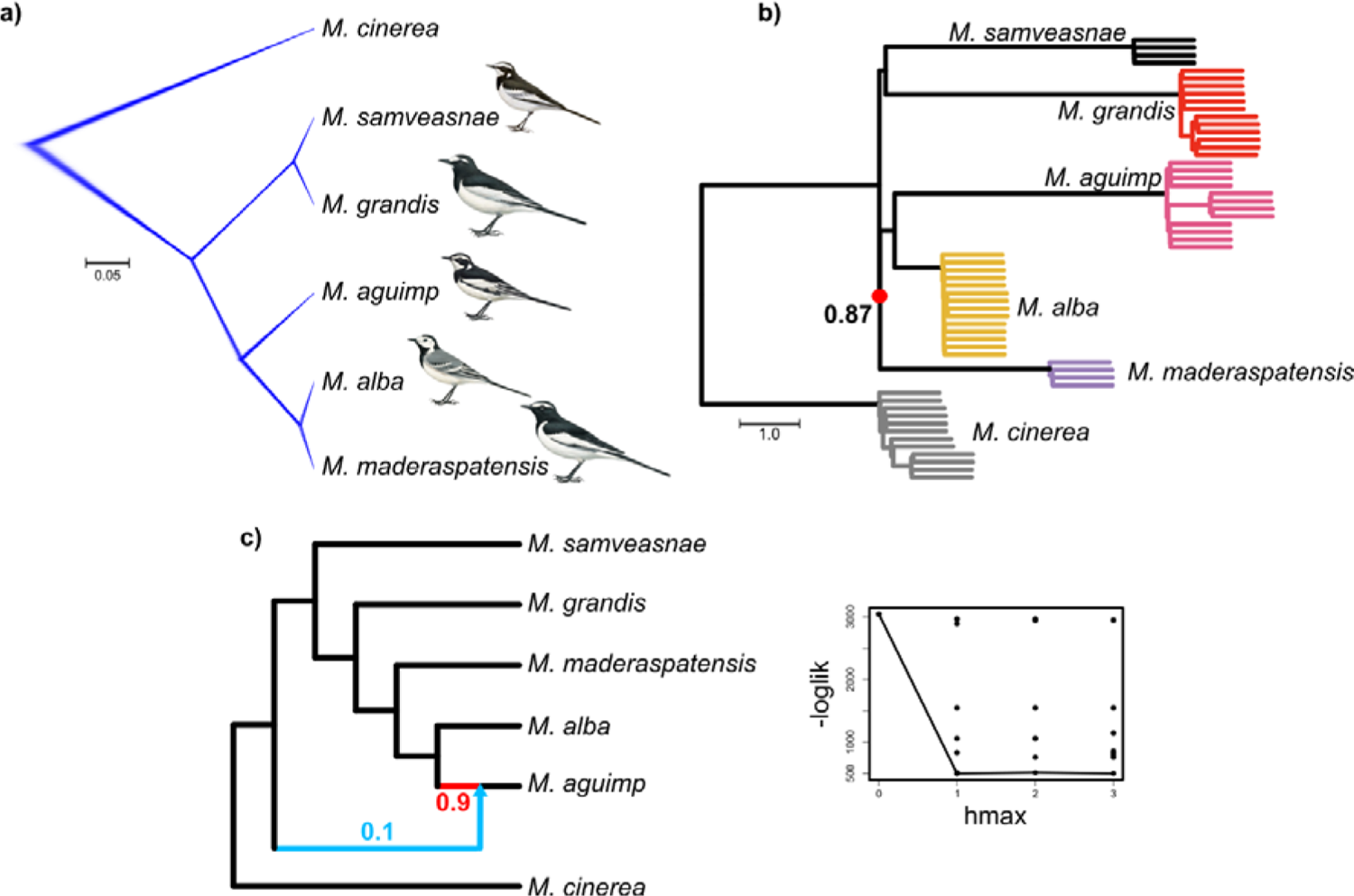
Results of phylogenetic analyses. A) cloudogram of the posterior distribution of species trees inferred in SNAPPER from 167,186 unlinked autosomal SNPs. Branch lengths represent the number of expected mutations per site. All nodes have posterior probability of 1.0. b) ASTRAL tree inferred from 9,490 trees calculated in 50 kb non-overlapping and non-adjacent sliding windows. Branch lengths represent coalescence units. Local posterior probabilities are indicated when < 1.0. c) Best phylogenetic network inferred in SnaQ! from the same set of SNPs as in the SNAPPER analysis. The red edge and blue arrow show the major and minor edges, respectively, of the inferred hybridization event, and the associated numbers are their inheritance probabilities (γ). The graph on the right shows the change in loglik when increasing h_max_, i.e., the maximum number of reticulations (line = loglik of the best model for each h_max_).

**Figure 3.**
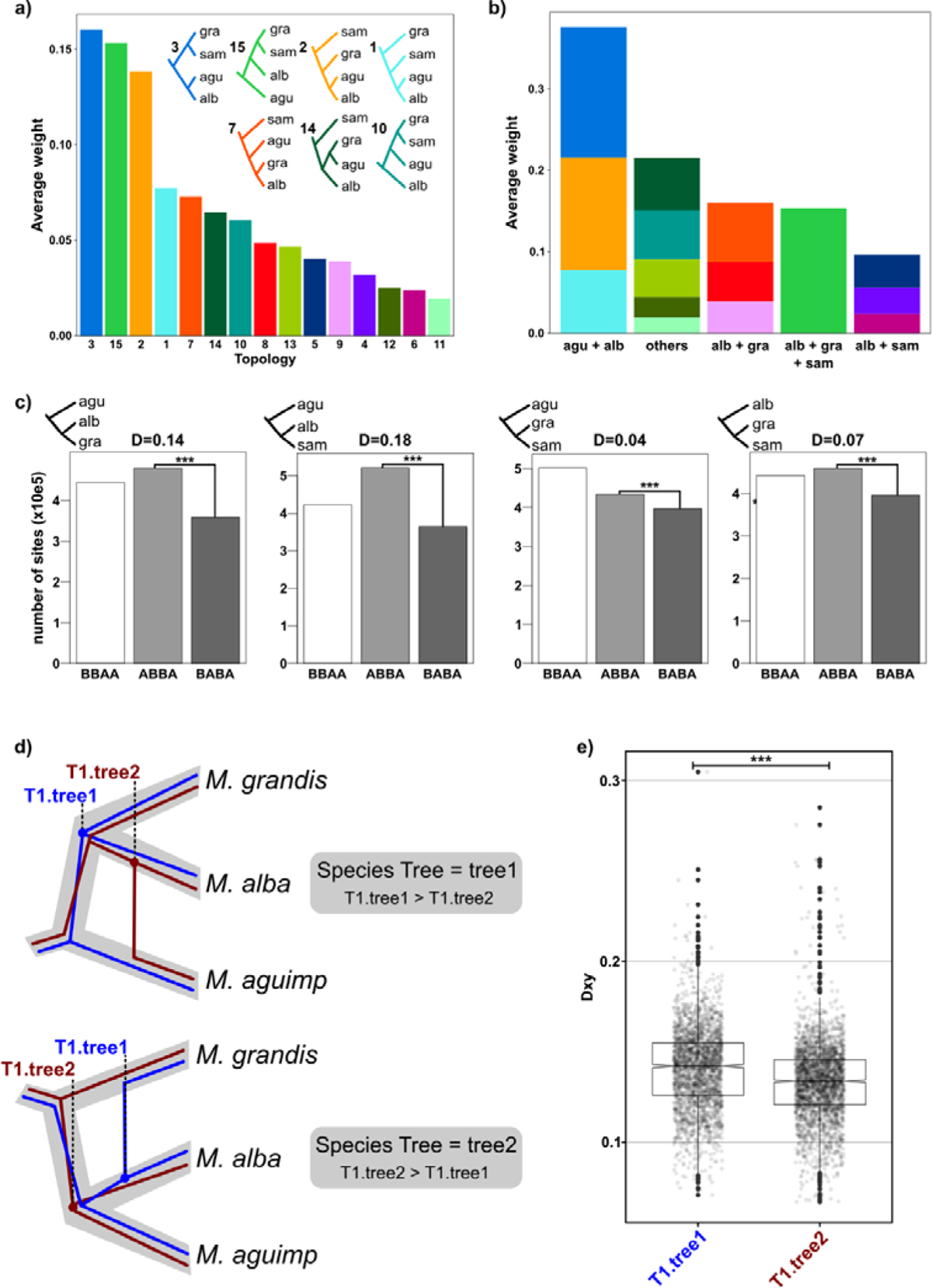
Variation of the phylogenetic signal across the genome. a) Average weights of the 15 alternative topologies (rooted with M. cinerea) calculated from trees in autosomal non-overlapping 50 kb sliding windows. The seven topologies with an average weight > 0.05 are shown (the outgroup was removed for improved graphical clarity). b) The same results with the 15 topologies aggregated in five categories depending on whether specific nodes are supported. c) Results of ABBA-BABA tests calculated from all autosomal SNPs, with top-left trees indicating the trio tested (P1–P3 from bottom to top, outgroup removed). agu = M. aguimp, alba = M. alba, gra = M. grandis, sam = M. samveasnae. d) Illustration of the “minimum height criterion” used to identify the species tree (i.e., the topology representing the history of speciation events) from introgression events. The gray shade represents the species tree, whereas the blue and red trees are gene trees supporting either M. alba + M. grandis or M. aguimp + M. alba, respectively. e) Height of the first split (approximated using D_XY_) in 50 kb sliding windows supporting either M. alba + M. grandis (T1.tree1) or M. aguimp + M. alba (T1.tree2).

In a similar fashion, we tested whether *M. alba* shares a more recent common ancestor with *grandis* or *grandis* + *samveasnae*. We considered the 50 kb sliding windows identified in the previous step as supporting the “non-introgressed” relationships, and compared the Dxy between *alba* and *grandis*, *alba* and *samveasnae* and *grandis* and *samveasnae* in these windows using the same test as before. Under a species tree where *alba* is sister to *grandis* + *samveasnae,* we expect that Dxy(*alba/grandis*) = Dxy(*alba/samveasnae*) > Dxy(*grandis/samveasnae*).

## 3 Results

### 3.1 Phylogenetic relationships based on plumage characters

A NJ analysis of 16 plumage characters scored in breeding males (Fig. 1b, Fig. S1) recovered both *M. aguimp*/*M. samveasnae* (pairwise distance=2) and *M. grandis*/*M. maderaspatensis* (pairwise distance=4) as sister species, highlighting their respective similarity. In contrast, the subspecies of *M. alba* did not form a monophyletic group but were scattered across the tree (pairwise distances ranging 2–8).

### 3.2 Multispecies Coalescent species tree and network

As a first approach to estimate phylogenetic relationships within the black-and-white wagtails while accounting for ILS and ILS+introgression, respectively, we used three Multispecies Coalescent models. When performing species tree estimation with SNAPPER based on the *snapper_5sp* dataset, individual runs failed to reach convergence. However, combining results from three independent runs improved the results: although several parameters still had ESS <200, posterior traces did not show clear trends. Despite suboptimal convergence, all three runs individually, as well as combined, yielded a single, fully supported topology (i.e., all nodes with Posterior Probability [PP] = 1.0, Fig. 2a). In this tree, *M. samveasnae/M. grandis* and *M. maderaspatensis*/*M. alba* form two pairs of sister species, and *M. aguimp* is sister to the latter. When excluding *M. maderaspatensis* this analysis converged properly (Fig. S2) on the same topology (i.e., *M. samveasnae*/*M. grandis* and *M. aguimp*/*M.alba* formed reciprocally monophyletic groups). The analysis based on Z chromosome variants yielded the same topology, although with lower support (Fig. S3a). The *ASTRAL_5sp* and *ASTRAL_4sp* analysis yielded similar topologies to their SNAPPER counterparts (Fig. 2b, Fig. S4), although in the *5sp* analysis the positions of *M. aguimp* and *M. maderaspatensis* were swapped, but this node received comparatively low support (PP = 0.87). Internal branches of the ingroup were short in all analyses, especially the ancestral branch of *M. aguimp, M. alba* and *M. maderaspatensis* in the ASTRAL *5sp* analysis (Fig. 2b).

In the multispecies coalescent network analyses of both the *snapper_5sp* (Fig. 2c) and *snapper_4sp* (Fig. S5) datasets, the loglik score decreased noticeably when introducing one reticulation, but remained stationary when adding two or three reticulations. Even when setting *h_max_* to 2 or 3, only one reticulation was recovered. Thus, we considered the network with a single reticulation and lowest loglik as best fitting our data. Both datasets yielded a network with a reticulation in the ancestry of *M. aguimp*. The two trees induced by the network placed *M. aguimp* either as sister to *M. alba* (major edge in both cases, inheritance probability [γ] = 0.90 and 0.65 in the *snapper_5sp* and *snapper_4sp* datasets, respectively) or as sister to all other ingroup species (minor edge, γ=0.10 and 0.35, respectively). In other words, the ancestral branch of *M. alba* and *M. aguimp* contributed between 65% and 90% to the genomic diversity of *M. aguimp*.

### 3.3 Genome-wide patterns of phylogenetic conflicts

Topology weighting revealed substantial levels of phylogenetic conflicts, as all 15 possible topologies were recovered (Fig. 3a). Among those, three were dominant, with an average weighting of 0.13 – 0.16 each: 1) the topology supported by SNAPPER, i.e*. M. alba* + *M. aguimp* and *M. samveasnae* + *M. grandis* (topology 3); 2) a topology placing *alba* as sister species to *M. samveasnae* + *M. grandis*, with *M. aguimp* as sister to the three others (topology 15); and 3) a topology placing *M. alba* and *M. aguimp* as sister species, with *M. grandis* sister to these and *M. samveasnae* sister to the three others (topology 2). Four alternative topologies received average weighting >0.05, while the remaining topologies received between 0.02 and 0.05. Clustering these topologies in five categories depending on the species they support as sisters (Fig. 3b) showed that topologies supporting *M. aguimp* + *alba* were the most frequently recovered (cumulative average weight = 0.38). Topologies supporting either *M. alba* + *M. grandis* or *M. alba* + (*M. grandis* + *M. samveasnae*) had weights slightly above 0.15, while those supporting *M. alba* + *M. samveasnae* had an average weight of 0.09. Finally, five topologies supporting alternative branching had a cumulative average weight of 0.20.

Topology weighting based on the Z chromosome yielded overall similar results (Fig. S3b–d), with all 15 topologies represented. As in the analysis of autosomal data, the topologies placing *M. aguimp* and *M. alba* as sisters were the most common (cumulative average weight = 0.51). However, topologies placing *M. alba* and *M. grandis* as sisters were the second most commonly supported topology on the Z chromosome (cumulative average weight = 0.26). Differences between the autosomal and Z chromosome results mostly concerned two topologies: the topology (*M. aguimp*,(*M. alba*,(*M. samveasnae*,*M. grandis*))) was more represented in the autosomal trees (topology 15, average weights of 0.16 and 0.06, respectively), while the topology (*M. samveasnae*,(*M. aguimp*,(*M. alba*, *M. grandis*))) was more common in the Z chromosome trees (topology 7, average weights of 0.07 and 0.16, respectively).

*D* statistics yielded significant signals for introgression in the four tested species trios (p-value=0, Z > 9; Fig. 3c; Table S3). The highest *D* statistics were found for the two trios with P1= either *M. samveasnaes* or *M. grandis*, P2=*M. alba* and P3=*M. aguimp* (*D*=0.18 and 0.14, respectively). In the two remaining trios, with P1=*M. samveasnae*, P2=*M. grandis* and P3=either *M. aguimp* or *M. alba*, *D* statistics were 0.04 and 0.07, respectively. The respective counts of BBAA, ABBA and BABA sites varied widely depending on the trio considered (Fig. 3c; Table S3), and ABBA sites were the most common in three of them. Overall, the differences in counts across the three site categories were low, as the least common site pattern represented 70–86% of the most common one, indicating high levels of ILS. *f_b_* statistics calculated based on the topology (*M. aguimp*, (*M. alba*, (*M. samveasnae*, *M. grandis*))) supported three introgression events (Fig. 4a): between the branches of *M. aguimp* and *M. alba* (*f_b_*=0.13, Z=39.44), between the branches of *M. alba* and *M. grandis* (*f_b_*=0.16, Z=13.28), and between the branches of *M. aguimp* and *M. grandis* (*f_b_*=0.03, Z=9.44). When calculated based on the topology ((*M. aguimp, M. alba*), (*M. samveasnae, M. grandis*)), *f_b_*statistics also supported the two latter introgression events (Fig. S6), plus one between the branches of *M. samveasnae* and *M. alba*.

**Figure 4.**
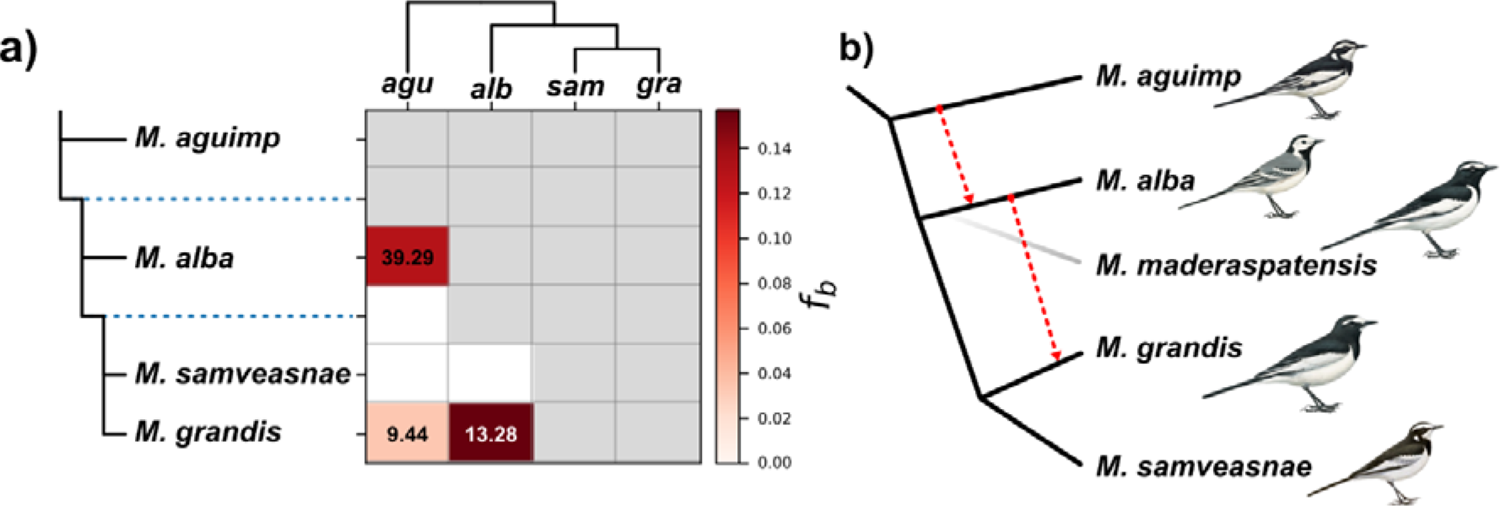
Proposed phylogenetic hypothesis. a) f_b_ statistic matrix. Numbers in the cells indicate the Z scores of the inferred introgression events. agu=aguimp; alb=alba; sam=samveasnae; gra=grandis. b) Proposed phylogenetic network for the black and white wagtails, based on the result of the various analyses presented here. The black lines represent our preferred species tree, and red arrows introgression events. The shaded gray branch of M. maderaspatensis highlights its uncertain position. Paintings courtesy Bill Zetterström (from Alström & Mild 2003).

### 3.4 Phylogenetic position of M. aguimp and M. alba

We investigated whether the sister relationship between *M. alba* and *M. aguimp* could be the result of introgression by comparing the Dxy between *M. alba* and *M. grandis* or between *M. alba* and *M. aguimp,* respectively, in genomic regions supporting either pair as monophyletic (i.e., supporting either “tree1” or “tree2” in Fig. 3d). We found that the D_XY_ between *M. grandis* and *M. alba* in regions supporting “tree1” (T1.tree1) was significantly higher than that between *M. aguimp* and *M. alba* in regions supporting “tree2” (T1.tree2; p-value < 2.2e-10). In other words, coalescence events defining the sister relationship between *M. aguimp* and *M. alba* are more recent than those between *M. grandis* and *M. alba.* This pattern is consistent with a sister relationship between the *M. grandis* and *M. alba* lineages, and later introgression between the latter and the *M. aguimp* lineage, meaning that “tree1” best represents the species tree. Based on this conclusion, we applied a similar approach to genomic regions identified as non-introgressed (i.e., supporting “tree1”) in the previous step, to determine whether *M. alba* is sister species to *M. grandis* or to *M. grandis* + *M. samveasnae*. In these regions, we found that the Dxy between *M. alba* and *M. grandis* did not differ from that between *M. alba* and *M. samveasnae* (Fig. S7, p-value = 0.23), consistent with the latter hypothesis. However, we also found that D_XY_ between *M. grandis* and *M. samveasnae* was significantly higher than that between the previous two pairs (p-value = 2.8e-08 and 2.3e-05, respectively).

## 4 Discussion

### Incomplete Lineage Sorting and ancient introgression obscure the relationships among black-and-white wagtail species

Our analyses of genome-wide polymorphisms confirm that relationships among the black-and-white wagtails are obscured by strong phylogenetic heterogeneity across the genome. All 15 possible topologies are represented, and even the three dominant topologies are in relatively low proportions (13–16%). This pattern, alongside the very short internal branches in phylogenetic trees, high levels of shared derived alleles between species, and their recent divergence (ca 0.7–2 Ma; Harris et al., 2018, Alström et al., 2015) point towards ILS as a major driver of phylogenetic conflict. Phylogenetic methods accounting for ILS yielded strong support for a topology dividing the group into two groups of species: *M. grandis* + *M. samveasnae* and *M. aguimp* + *M. maderaspatensis* + *M. alba*, respectively. However, we also recovered strong signals of introgression from both phylogenetic networks and *D* statistics analyses. Under these circumstances, determining which topology represents the species tree (a term used here to refer to the phylogenetic tree representing the history of speciation events) is not straightforward: the inferred network does not explicitly inform on which of its induced trees represents the species tree (as opposed to introgression events; Hibbins & Hahn, 2022b), while interpreting *D* statistics requires an explicit assumption of the species tree. Using a “minimum node height” criterion, we found that the most recent split in gene trees supporting *M. alba* as sister to *M. aguimp* was shallower than in gene trees supporting *M. alba* as sister to *M. grandis*. This result supports that the sister relationship of *M. aguimp* and *M. alba*, which is predominantly found across the genome, is influenced by ancient introgression. Other cases where the dominant gene tree topology is attributed to introgression include *Phylloscopus* leaf-warblers (Zhang et al., 2021), *Lonchura* munias (Stryjewski & Sorenson, 2017) and *Anopheles* mosquitoes (Fontaine et al. 2015). It should be noted, however, that this does not mean that all regions supporting *M. alba* and *M. aguimp* as sister species are introgressed. Part of these gene trees have likely been generated by ILS, as indicated by the difference between their average weight across the genome and the *f_b_* statistic for this introgression event (0.39 vs. 0.13, respectively). Furthermore, when sorting is incomplete, introgression might reintroduce standing variation that can subsequently be differentially fixed, meaning that gene trees at loci introgressed between *M. aguimp* and *M. alba* will not necessarily support them as sister species. Considering this, topologies alone do not provide enough information to identify introgressed regions, and by extension identify the species tree. Simulations by Hibbins & Hahn (2022b) show that the most recent split in introgressed gene trees is not necessarily shallower than in non-introgressed regions, especially in cases where speciation and introgression events closely follow each other, and both the rates of introgression and background discordance (i.e., ILS) are high. However, the estimated rate of introgression necessary for introgressed gene trees to appear deeper than those representing the species tree (≥20%, except for rates of ILS ≥60%; Hibbins and Hahn 2022b) remains above the 13% of introgression between *M. aguimp* and *M. alba* estimated by the *f_b_* statistic. *D* and *f_b_* statistics calculated based on the “tree1” topology are also more parsimonious, as they infer fewer introgression events. Hence, we favor a species tree where *M. aguimp* is sister to the other four black-and-white wagtail species (Fig. 4b).

*Motacilla grandis* was recovered as sister to *M. samveasnae* in all but one analysis. Considering the comparatively high proportion of gene trees placing *M. alba* and *M. grandis* as sister species, it is possible that this variation denotes a second introgression event. It is relevant to note that inferences of phylogenetic networks in SNaQ! are limited to level-1 networks, i.e., a maximum of one reticulation can occur along a given branch of the tree (Solís-Lemus and Ané 2016), because higher-level networks are not identifiable from sequence or gene tree data under multispecies coalescent network models (Pardi and Scornavacca 2015). Hence, SNaQ! cannot accurately reconstruct the species network if several introgression events occurred along the ancestral branch of *M. alba*. We favor a species tree where *M. alba* is sister to a monophyletic *M. grandis* + *M. samveasnae*, with an introgression event between the *M. alba* and *M. grandis* lineages, as supported by *D* and *f_b_* statistics.

This is indirectly supported by the fact that both statistics also inferred an introgression event between *M. aguimp* and *M. grandis*. Indeed, while introgressive hybridization between the branches of *M. aguimp* and *M. grandis* does not seem likely because of their wide geographical separation (Fig. 1), this pattern of allele sharing could be explained if introgression from *M. aguimp* into *M. alba* was followed by introgression from the latter into *M. grandis*, allowing alleles to “travel” from *M. aguimp* to *M. grandis* (as found in Darwin’s finches; Grant and Grant 2020). We attempted to confirm this result using the “minimum node height” criterion, but the results were inconclusive, possibly because of differences in the demographic histories of the compared species. It is quite surprising that the introgressed topology placing *M. alba* and *M. grandis* as sister species is much more common on the Z chromosome compared to the autosomes. It has been suggested that sex chromosomes are less porous to introgression, and therefore more likely to support the species tree topology (Qvarnström and Bailey 2009; Fontaine et al. 2015), but this does not seem to be the case in the black-and-white wagtails.

Overall, our results show that wagtail genomes are mosaics of regions supporting variable genealogies, because of ILS and inter-specific introgression. Under these circumstances, identifying the species tree is far from trivial, but we reconstructed the species network (i.e., a network representing both speciation and introgression events) with confidence by combining complementary approaches to infer phylogenies and introgression (Fig. 4b). Identifying which of the topologies induced by the network represents the species tree remains an outstanding challenge, especially when the most common topology is influenced by introgression. Further studies of the characteristics of non-introgressed vs. introgressed gene trees will likely help defining separation criteria more precisely (Hibbins and Hahn 2022b). However, it should be noted that, in this case, uncertainty on the species tree topology does not prevent inferences of patterns of phenotypic evolution. Resolving the species network, or part of it, yields important information on the presence of parallel phenotypic evolution, and the contribution of introgression to this pattern.

### Introgression does not underlie parallel plumage evolution in the “black-and-white” wagtails

Based on our analyses of genomic data, we propose an evolutionary scenario where *M. aguimp* diverged first from the other black-and-white wagtail species, followed by a split between *M. alba* and *M. grandis* + *M. samveasnae* (Fig. 4b). The position of *M. maderaspatensis* is less certain, but most of our analyses place it as sister to *M. alba* (to the exception of the ASTRAL analysis where *M. maderaspatensis* is sister to *M. alba* and *M. aguimp*). Furthermore, we evidence two instances of introgressive hybridization, firstly between *M. aguimp* and *M. alba,* and secondly between *M. alba* and *M. grandis* (Fig. 4b).

In contrast, analyses of plumage characteristics support that *M. aguimp* + *M. samveasnae* and *M. grandis* + *M. maderaspatensis* are very similar to each other, and distinct from *M. alba*. Hence, our phylogenomic results confirm that similar plumages evolved in non-sister lineages. The remaining uncertainty regarding which topology represents the species tree does not invalidate this conclusion, as none of the topologies induced by the network place similar-looking species (*M. aguimp* + *M. samveasnae* and *M. grandis* + *M. maderaspatensis*) together. Indeed, while introgressive hybridization has occurred, inferred reticulations do not connect lineages with similar phenotypes, contradicting the hypothesis that parallel plumage evolution is caused by horizontal transfer of alleles. It is possible, however, that the variants underlying some key plumage traits are older than the species themselves and fixed independently in non-sister lineages (i.e., because of ILS). ILS has been shown to be responsible for parallel phenotypic evolution in several cases (e.g., Pease et al. 2016; Feng et al. 2022), including plumage characters in the passerine genus *Oenanthe* (Alaei Kakhki et al. 2023). However, lineage-specific *de novo* mutations should also be considered as a plausible explanation: black coloration in pigs evolved following new mutations in the MC1R gene emerging after domestication (<10,000 years ago, Fang et al. 2009), a short timescale compared to the *Motacilla* radiation. It is possible that both of these processes contribute to plumage evolution in the black-and-white wagtails (as found in the genus *Oenanthe*; Alaei Kakhki et al. 2023), as several loci are involved (Semenov et al. 2021). Future efforts will be directed towards identifying genomic variants associated with plumage traits, to explicitly test these hypotheses.

Another fascinating aspect of the black-and-white wagtails is the high phenotypic diversity of *M. alba* compared to its closest relatives. It might be relevant to link that pattern with the two introgression events we recovered in the ancestry of *M. alba*. Indeed, introgression can act as a driver of diversification, by generating new combinations of variants, and potentially novel phenotypes (Stryjewski and Sorenson 2017; Marques et al. 2019; Svardal et al. 2020). Such enriched genetic background, combined with biogeographic events (e.g., populations fragmentation linked to glacial cycles; Milá et al. 2007a, 2007b; Weir et al. 2016), and strong selective pressures could have led *M. alba* to evolve such geographic variations in plumage phenotypes. In the light of our results, future investigations will focus on the role of introgressed variants in plumage divergence across subspecies of *M. alba*.

## Conclusion

Our results build on the vast literature showing that species tree inference in the face of gene flow and ILS is an outstanding challenge. Especially, we show that the most common gene tree in the “black-and-white” wagtails might be the result of introgression, a pattern that has been rarely described. However, we demonstrate that a bifurcating species tree is not a prerequisite to investigate patterns of parallel phenotypic evolution. In the presence of introgression, inferring the species network will always be easier than reconstructing the species tree. Even if the latter cannot be determined with confidence, conclusions can be drawn from whether one (or several) of the trees induced by the network place species with similar phenotypes as sisters. It should be noted, however, that the presence of introgression between similar phenotypes does not guarantee that this process underlies phenotypic evolution. Ultimately, a detailed understanding of the evolution of phenotypes requires to uncover their genomic basis. Likewise, identifying the species tree is useful to understand the speciation history.

## Acknowledgments

We are grateful to Martin Stervander for his help with the snapper analyses. P.A. was supported by the Swedish Research Council (2015-04402 and 2019-04486), Jornvall Foundation and Carl Trygger Foundation (CTS 20:6). LA is supported by Swedish Research Council (2017-02907) and Knut and Alice Wallenberg Foundation (KAW 2016.0361). FL was supported by the Second Tibetan Plateau Scientific Expedition and Research (STEP) program (Grant No. 2019QZKK0304-02).

## Authors contributions

PA, LA, EDE and LR conceived the study. TS, YL and FL provided samples. RH and MI prepared libraries. EDE performed the variant calling and preliminary explorations of the data. PA scored the plumage characteristics. LR performed additional filtering of the genomic data and analyses. LR wrote the manuscript with input from EDE and PA. All co-authors contributed to improve the manuscript.

## Data availability statement

The data files used to perform the analyses in this manuscript, as well as the supplementary materials, are available at <u>##### link to dryad repository to be added upon submission #####</u>.

## References

Alaei Kakhki N., Schweizer M., Lutgen D., Bowie R.C.K., Shirihai H., Suh A., Schielzeth H., Burri R. 2023. A Phylogenomic Assessment of Processes Underpinning Convergent Evolution in Open-Habitat Chats. Mol. Biol. Evol. 40:msac278.

Alström P., Jønsson K.A., Fjeldså J., Ödeen A., Ericson P.G.P., Irestedt M. 2015. Dramatic niche shifts and morphological change in two insular bird species. R. Soc. Open Sci. 2:140364.

Alström P., Mild K. 2003. Pipits and Wagtails of Europe, Asia and North America. London, UK: Helm/A&C Black.

Astudillo-Clavijo V., Stiassny M.L.J., Ilves K.L., Musilova Z., Salzburger W., López-Fernández H. 2022. Exon-based Phylogenomics and the Relationships of African Cichlid Fishes: Tackling the Challenges of Reconstructing Phylogenies with Repeated Rapid Radiations. Syst. Biol.:syac051.

Baack E.J., Rieseberg L.H. 2007. A genomic view of introgression and hybrid speciation. Curr. Opin. Genet. Dev. 17:513–518.

Beckman E.J., Benham P.M., Cheviron Z.A., Witt C.C. 2018. Detecting introgression despite phylogenetic uncertainty: The case of the South American siskins. Mol. Ecol. 27:4350–4367.

Bouckaert R.R. 2010. DensiTree: making sense of sets of phylogenetic trees. Bioinformatics. 26:1372–1373.

Cai L., Xi Z., Lemmon E.M., Lemmon A.R., Mast A., Buddenhagen C.E., Liu L., Davis C.C. 2021. The Perfect Storm: Gene Tree Estimation Error, Incomplete Lineage Sorting, and Ancient Gene Flow Explain the Most Recalcitrant Ancient Angiosperm Clade, Malpighiales. Syst. Biol. 70:491–507.

Chen S., Zhou Y., Chen Y., Gu J. 2018. fastp: an ultra-fast all-in-one FASTQ preprocessor. Bioinformatics. 34:i884–i890.

Chen W.-C. 2011. Overlapping codon model, phylogenetic clustering, and alternative partial expectation conditional maximization algorithm. Doctoral dissertation, Iowa State University.

Dasmahapatra K.K., Walters J.R., Briscoe A.D., Davey J.W., Whibley A., Nadeau N.J., Zimin A.V., Hughes D.S.T., Ferguson L.C., Martin S.H., Salazar C., Lewis J.J., Adler S., Ahn S.-J., Baker D.A., Baxter S.W., Chamberlain N.L., Chauhan R., Counterman B.A., Dalmay T., Gilbert L.E., Gordon K., Heckel D.G., Hines H.M., Hoff K.J., Holland P.W.H., Jacquin-Joly E., Jiggins F.M., Jones R.T., Kapan D.D., Kersey P., Lamas G., Lawson D., Mapleson D., Maroja L.S., Martin A., Moxon S., Palmer W.J., Papa R., Papanicolaou A., Pauchet Y., Ray D.A., Rosser N., Salzberg S.L., Supple M.A., Surridge A., Tenger-Trolander A., Vogel H., Wilkinson P.A., Wilson D., Yorke J.A., Yuan F., Balmuth A.L., Eland C., Gharbi K., Thomson M., Gibbs R.A., Han Y., Jayaseelan J.C., Kovar C., Mathew T., Muzny D.M., Ongeri F., Pu L.-L., Qu J., Thornton R.L., Worley K.C., Wu Y.-Q., Linares M., Blaxter M.L., ffrench-Constant R.H., Joron M., Kronforst M.R., Mullen S.P., Reed R.D., Scherer S.E., Richards S., Mallet J., Owen McMillan W., Jiggins C.D., The Heliconius Genome Consortium. 2012. Butterfly genome reveals promiscuous exchange of mimicry adaptations among species. Nature. 487:94–98.

Degnan J.H., Rosenberg N.A. 2009. Gene tree discordance, phylogenetic inference and the multispecies coalescent. Trends Ecol. Evol. 24:332–340.

Delaneau O., Zagury J.-F., Robinson M.R., Marchini J.L., Dermitzakis E.T. 2019. Accurate, scalable and integrative haplotype estimation. Nat. Commun. 10:5436.

Drovetski S.V., Reeves A.B., Red’kin Y.A., Fadeev I.V., Koblik E.A., Sotnikov V.N., Voelker G. 2018. Multi-locus reassessment of a striking discord between mtDNA gene trees and taxonomy across two congeneric species complexes. Mol. Phylogenet. Evol. 120:43–52.

Durand E.Y., Patterson N., Reich D., Slatkin M. 2011. Testing for Ancient Admixture between Closely Related Populations. Mol. Biol. Evol. 28:2239–2252.

Elmer K.R., Meyer A. 2011. Adaptation in the age of ecological genomics: insights from parallelism and convergence. Trends Ecol. Evol. 26:298–306.

Enciso-Romero J., Pardo-Díaz C., Martin S.H., Arias C.F., Linares M., McMillan W.O., Jiggins C.D., Salazar C. 2017. Evolution of novel mimicry rings facilitated by adaptive introgression in tropical butterflies. Mol. Ecol. 26:5160–5172.

Fang M., Larson G., Ribeiro H.S., Li N., Andersson L. 2009. Contrasting Mode of Evolution at a Coat Color Locus in Wild and Domestic Pigs. PLOS Genet. 5:e1000341.

Feng S., Bai M., Rivas-González I., Li C., Liu S., Tong Y., Yang H., Chen G., Xie D., Sears K.E., Franco L.M., Gaitan-Espitia J.D., Nespolo R.F., Johnson W.E., Yang H., Brandies P.A., Hogg C.J., Belov K., Renfree M.B., Helgen K.M., Boomsma J.J., Schierup M.H., Zhang G. 2022. Incomplete lineage sorting and phenotypic evolution in marsupials. Cell. 185:1646–1660.e18.

Fontaine M.C., Pease J.B., Steele A., Waterhouse R.M., Neafsey D.E., Sharakhov I.V., Jiang X., Hall A.B., Catteruccia F., Kakani E., Mitchell S.N., Wu Y.-C., Smith H.A., Love R.R., Lawniczak M.K., Slotman M.A., Emrich S.J., Hahn M.W., Besansky N.J. 2015. Extensive introgression in a malaria vector species complex revealed by phylogenomics. Science. 347:1258524.

Forsythe E.S., Nelson A.D.L., Beilstein M.A. 2020. Biased Gene Retention in the Face of Introgression Obscures Species Relationships. Genome Biol. Evol. 12:1646–1663.

Freed D., Aldana R., Weber J.A., Edwards J.S. 2017. The Sentieon Genomics Tools - A fast and accurate solution to variant calling from next-generation sequence data. BioRxiv, https://doi.org/10.1101/115717.

Giarla T.C., Esselstyn J.A. 2015. The Challenges of Resolving a Rapid, Recent Radiation: Empirical and Simulated Phylogenomics of Philippine Shrews. Syst. Biol. 64:727–740.

Grant P.R., Grant B.R. 2020. Triad hybridization via a conduit species. Proc. Natl. Acad. Sci. 117:7888–7896.

Green R.E., Krause J., Briggs A.W., Maricic T., Stenzel U., Kircher M., Patterson N., Li H., Zhai W., Fritz M.H.-Y., Hansen N.F., Durand E.Y., Malaspinas A.-S., Jensen J.D., Marques-Bonet T., Alkan C., Prüfer K., Meyer M., Burbano H.A., Good J.M., Schultz R., Aximu-Petri A., Butthof A., Höber B., Höffner B., Siegemund M., Weihmann A., Nusbaum C., Lander E.S., Russ C., Novod N., Affourtit J., Egholm M., Verna C., Rudan P., Brajkovic D., Kucan Ž., Gušic I., Doronichev V.B., Golovanova L.V., Lalueza-Fox C., de la Rasilla M., Fortea J., Rosas A., Schmitz R.W., Johnson P.L.F., Eichler E.E., Falush D., Birney E., Mullikin J.C., Slatkin M., Nielsen R., Kelso J., Lachmann M., Reich D., Pääbo S. 2010. A Draft Sequence of the Neandertal Genome. Science. 328:710–722.

Han F., Lamichhaney S., Grant B.R., Grant P.R., Andersson L., Webster M.T. 2017. Gene flow, ancient polymorphism, and ecological adaptation shape the genomic landscape of divergence among Darwin’s finches. Genome Res. 27:1004–1015.

Harris R.B., Alström P., Ödeen A., Leaché A.D. 2018. Discordance between genomic divergence and phenotypic variation in a rapidly evolving avian genus (*Motacilla*). Mol. Phylogenet. Evol. 120:183–195.

Hibbins M.S., Hahn M.W. 2022a. Phylogenomic approaches to detecting and characterizing introgression. Genetics. 220:iyab173.

Hibbins M.S., Hahn M.W. 2022b. Distinguishing between histories of speciation and introgression using genomic data. BioRxiv, https://doi.org/10.1101/2022.09.07.506990.

Lamichhaney S., Berglund J., Almén M.S., Maqbool K., Grabherr M., Martinez-Barrio A., Promerová M., Rubin C.-J., Wang C., Zamani N., Grant B.R., Grant P.R., Webster M.T., Andersson L. 2015. Evolution of Darwin’s finches and their beaks revealed by genome sequencing. Nature. 518:371–375.

Li G., Figueiró H.V., Eizirik E., Murphy W.J. 2019. Recombination-Aware Phylogenomics Reveals the Structured Genomic Landscape of Hybridizing Cat Species. Mol. Biol. Evol. 36:2111– 2126.

Maddison W.P., Knowles L.L. 2006. Inferring Phylogeny Despite Incomplete Lineage Sorting. Syst. Biol. 55:21–30.

Malinsky M., Matschiner M., Svardal H. 2021. Dsuite - Fast D-statistics and related admixture evidence from VCF files. Mol. Ecol. Resour. 21:584–595.

Marques D.A., Meier J.I., Seehausen O. 2019. A Combinatorial View on Speciation and Adaptive Radiation. Trends Ecol. Evol. 34:531–544.

Martin M., Patterson M., Garg S., Fischer S.O., Pisanti N., Klau G.W., Schöenhuth A., Marschall T. 2016. WhatsHap: fast and accurate read-based phasing. BioRxiv, https://doi.org/10.1101/085050.

Martin S.H., Van Belleghem S.M. 2017. Exploring Evolutionary Relationships Across the Genome Using Topology Weighting. Genetics. 206:429–438.

McKenna A., Hanna M., Banks E., Sivachenko A., Cibulskis K., Kernytsky A., Garimella K., Altshuler D., Gabriel S., Daly M., DePristo M.A. 2010. The Genome Analysis Toolkit: A MapReduce framework for analyzing next-generation DNA sequencing data. Genome Res. 20:1297–1303.

Milá B., McCormack J.E., Castañeda G., Wayne R.K., Smith T.B. 2007a. Recent postglacial range expansion drives the rapid diversification of a songbird lineage in the genus Junco. Proc. R. Soc. B Biol. Sci. 274:2653–2660.

Milá B., Smith T.B., Wayne R.K. 2007b. Speciation and rapid phenotypic differentiation in the yellow-rumped warbler *Dendroica coronata* complex. Mol. Ecol. 16:159–173.

Mirarab S., Reaz R., Bayzid Md.S., Zimmermann T., Swenson M.S., Warnow T. 2014. ASTRAL: genome-scale coalescent-based species tree estimation. Bioinformatics. 30:i541–i548.

Nichols R. 2001. Gene trees and species trees are not the same. Trends Ecol. Evol. 16:358–364.

Olave M., Meyer A. 2020. Implementing Large Genomic Single Nucleotide Polymorphism Data Sets in Phylogenetic Network Reconstructions: A Case Study of Particularly Rapid Radiations of Cichlid Fish. Syst. Biol. 69:848–862.

Ortiz E.M. 2019. vcf2phylip v2.0: convert a VCF matrix into several matrix formats for phylogenetic analysis. https://doi.org/10.5281/zenodo.2540861.

Pardi F., Scornavacca C. 2015. Reconstructible Phylogenetic Networks: Do Not Distinguish the Indistinguishable. PLOS Comput. Biol. 11:e1004135.

Pease J.B. 2018. Why phylogenomic uncertainty enhances introgression analyses. Mol. Ecol. 27:4347–4349.

Pease J.B., Haak D.C., Hahn M.W., Moyle L.C. 2016. Phylogenomics Reveals Three Sources of Adaptive Variation during a Rapid Radiation. PLOS Biol. 14:e1002379.

Qvarnström A., Bailey R.I. 2009. Speciation through evolution of sex-linked genes. Heredity. 102:4–15.

Rambaut A., Drummond A.J., Xie D., Baele G., Suchard M.A. 2018. Posterior Summarization in Bayesian Phylogenetics Using Tracer 1.7. Syst. Biol. 67:901–904.

Rannala B., Yang Z. 2003. Bayes Estimation of Species Divergence Times and Ancestral Population Sizes Using DNA Sequences From Multiple Loci. Genetics. 164:1645–1656.

Rheindt F.E., Edwards S.V. 2011. Genetic Introgression: An Integral but Neglected Component of Speciation in Birds. The Auk. 128:620–632.

Sayyari E., Whitfield J.B., Mirarab S. 2017. Fragmentary Gene Sequences Negatively Impact Gene Tree and Species Tree Reconstruction. Mol. Biol. Evol. 34:3279–3291.

Seehausen O. 2004. Hybridization and adaptive radiation. Trends Ecol. Evol. 19:198–207.

Semenov G.A., Linck E., Enbody E.D., Harris R.B., Khaydarov D.R., Alström P., Andersson L., Taylor S.A. 2021. Asymmetric introgression reveals the genetic architecture of a plumage trait. Nat. Commun. 12:1019.

Solís-Lemus C., Ané C. 2016. Inferring Phylogenetic Networks with Maximum Pseudolikelihood under Incomplete Lineage Sorting. PLOS Genet. 12:e1005896.

Solís-Lemus C., Bastide P., Ané C. 2017. PhyloNetworks: A Package for Phylogenetic Networks. Mol. Biol. Evol. 34:3292–3298.

Stamatakis A. 2014. RAxML version 8: a tool for phylogenetic analysis and post-analysis of large phylogenies. Bioinformatics. 30:1312–1313.

Stern D.L. 2013. The genetic causes of convergent evolution. Nat. Rev. Genet. 14:751–764.

Stoltz M., Baeumer B., Bouckaert R., Fox C., Hiscott G., Bryant D. 2021. Bayesian Inference of Species Trees using Diffusion Models. Syst. Biol. 70:145–161.

Stryjewski K.F., Sorenson M.D. 2017. Mosaic genome evolution in a recent and rapid avian radiation. Nat. Ecol. Evol. 1:1912–1922.

Svardal H., Quah F.X., Malinsky M., Ngatunga B.P., Miska E.A., Salzburger W., Genner M.J., Turner G.F., Durbin R. 2020. Ancestral Hybridization Facilitated Species Diversification in the Lake Malawi Cichlid Fish Adaptive Radiation. Mol. Biol. Evol. 37:1100–1113.

Wallbank R.W.R., Baxter S.W., Pardo-Diaz C., Hanly J.J., Martin S.H., Mallet J., Dasmahapatra K.K., Salazar C., Joron M., Nadeau N., McMillan W.O., Jiggins C.D. 2016. Evolutionary Novelty in a Butterfly Wing Pattern through Enhancer Shuffling. PLOS Biol. 14:e1002353.

Weir J.T., Haddrath O., Robertson H.A., Colbourne R.M., Baker A.J. 2016. Explosive ice age diversification of kiwi. Proc. Natl. Acad. Sci. 113:E5580–E5587.

Zhang C., Rabiee M., Sayyari E., Mirarab S. 2018. ASTRAL-III: polynomial time species tree reconstruction from partially resolved gene trees. BMC Bioinformatics. 19:153.

Zhang D., Rheindt F.E., She H., Cheng Y., Song G., Jia C., Qu Y., Alström P., Lei F. 2021. Most Genomic Loci Misrepresent the Phylogeny of an Avian Radiation Because of Ancient Gene Flow. Syst. Biol. 70:961–975.

