## Supplementary materials for "Introgression underlies phylogenetic uncertainty but not parallel plumage evolution in a recent songbird radiation"

**Supplementary material SM1: Details on the plumage characters dataset**

The following characters were used to calculate distances between black-and-white wagtail taxa. All characters were scored in adult male breeding plumage. Because the difference between each character state is qualitative and not quantitative, we calculated pairwise distances by assigning a distance of 1 if two taxa have a different state, and 0 if they have the same state. *Motacilla cinerea* was included as an outgroup.

1 – **forehead:** 0 – dark; 1 – thin dark stripe; 2 – white; 3 – white, often with incomplete dark stripe

2 – **crown, nape:** 0 – black; 1 – greyish

3 – **lores:** 0 – dark; 1 – white

4 – **distinct, discrete supercilium, not meeting across forehead:** 0 – present; 1 – absent

5 – **ear-coverts:** 0 – black; 1 – white; 2 – black; black eyestripe, white subocular; 3 – black, white subocular; 4 – white, black eyestripe; 5 – rather uniformly greyish, sometimes with paler patch below/behind eye

6 – **side of neck:** 0 – white; 1 – black; 2 – greyish

7 – **throat:** 0 – white; 1 – black, white submoustachial; 2 – black, short white submoustachial; 3 – black lower, white upper ≥half; 4 – black, white submoustachial, small white upper; 5 – black; 6 – black, small white upper

8 – **dark pattern on breast:** 0 – absent; 1 – black crescent extending to ear-coverts; 2 – black extends to throat

9 – **mantle and scapulars:** 0 – black; 1 – grey

10 – **rump:** 0 – black; 1 – grey; 2 – green

11 – **uppertail-coverts:** 0 – black; 4 – green

12 – **breast-sides and flanks:** 0 – white; 1 – pale to medium grey wash (may be mainly concealed); 2 – white or black; 3 – yellow breast-sides, pale yellow or whitish flanks

13 – **belly:** 0 – white; 1 – yellow

14 – **undertail-coverts:** 0 – white; 1 – yellow

15 – **median coverts (fresh):** 0 – blackish bases, rather narrow brownish or greyish tips, not forming distinct contrasting pale wing-bar; 1 – blackish bases, fairly broad pale tips, forming distinct contrasting pale wing-bar; 2 – visible part appearing all or nearly all white; 3 – variable, ranging from blackish centres, fairly broad pale tips, forming distinct contrasting pale wing-bar, or visible part appearing all or nearly all white

16 – **greater coverts (fresh):** 0 – blackish centres, narrow greyish tips; 1 – blackish centres, fairly broad pale tips, forming distinct contrasting pale wing-bar; 2 – visible part appearing all or nearly all white (together with white tips to median coverts forming all or mainly white panel); 3 – variable, ranging from blackish centres, fairly broad pale tips, forming distinct contrasting pale wing-bar, or visible part appearing all or nearly all white, forming all or mainly white panel

**Table S1.** Plumage characters matrix.

| **Character**  **Taxa** | **1** | **2** | 3 | 4 | 5 | 6 | **7** | **8** | **9** | **10** | **11** | **12** | **13** | **14** | **15** | **16** |
| --- | --- | --- | --- | --- | --- | --- | --- | --- | --- | --- | --- | --- | --- | --- | --- | --- |
| *M. aguimp* | 1 | 0 | 0 | 0 | 0 | 0 | 0 | 1 | 0 | 0 | 0 | 2 | 0 | 0 | 2 | 2 |
| *M. a. alba* | 2 | 0 | 1 | 1 | 1 | 0 | 1 | 2 | 1 | 1 | 0 | 1 | 0 | 0 | 1 | 1 |
| *M. a. yarrellii* | 2 | 0 | 1 | 1 | 1 | 0 | 1 | 2 | 0 | 0 | 0 | 1 | 0 | 0 | 1 | 1 |
| *M. a. subpersonata* | 2 | 0 | 0 | 1 | 2 | 0 | 2 | 2 | 1 | 1 | 0 | 1 | 0 | 0 | 1 | 1 |
| *M. a. baicalensis* | 2 | 0 | 1 | 1 | 1 | 0 | 3 | 2 | 1 | 1 | 0 | 1 | 0 | 0 | 3 | 3 |
| *M. a. ocularis* | 2 | 0 | 0 | 1 | 4 | 0 | 1 | 2 | 1 | 1 | 0 | 1 | 0 | 0 | 3 | 3 |
| *M. a. lugens* | 2 | 0 | 0 | 1 | 4 | 0 | 4 | 2 | 0 | 0 | 0 | 1 | 0 | 0 | 2 | 2 |
| *M. a. leucopsis* | 2 | 0 | 1 | 1 | 1 | 0 | 3 | 2 | 0 | 0 | 0 | 0 | 0 | 0 | 2 | 2 |
| *M.a. alboides* | 2 | 0 | 1 | 1 | 3 | 1 | 5 | 2 | 0 | 0 | 0 | 1 | 0 | 0 | 2 | 2 |
| *M.a. personata* | 2 | 0 | 1 | 1 | 3 | 1 | 5 | 2 | 1 | 1 | 0 | 1 | 0 | 0 | 2 | 2 |
| *M. maderaspatensis* | 1 | 0 | 0 | 0 | 0 | 1 | 5 | 2 | 0 | 0 | 0 | 1 | 0 | 0 | 2 | 2 |
| *M. grandis* | 3 | 0 | 0 | 1 | 0 | 1 | 6 | 2 | 0 | 0 | 0 | 0 | 0 | 0 | 2 | 2 |
| *M. samveasnae* | 1 | 0 | 0 | 0 | 0 | 0 | 0 | 1 | 0 | 0 | 0 | 0 | 0 | 0 | 2 | 3 |
| *M. cinerea* | 0 | 1 | 0 | 0 | 5 | 2 | 1 | 0 | 1 | 2 | 1 | 3 | 1 | 1 | 0 | 0 |

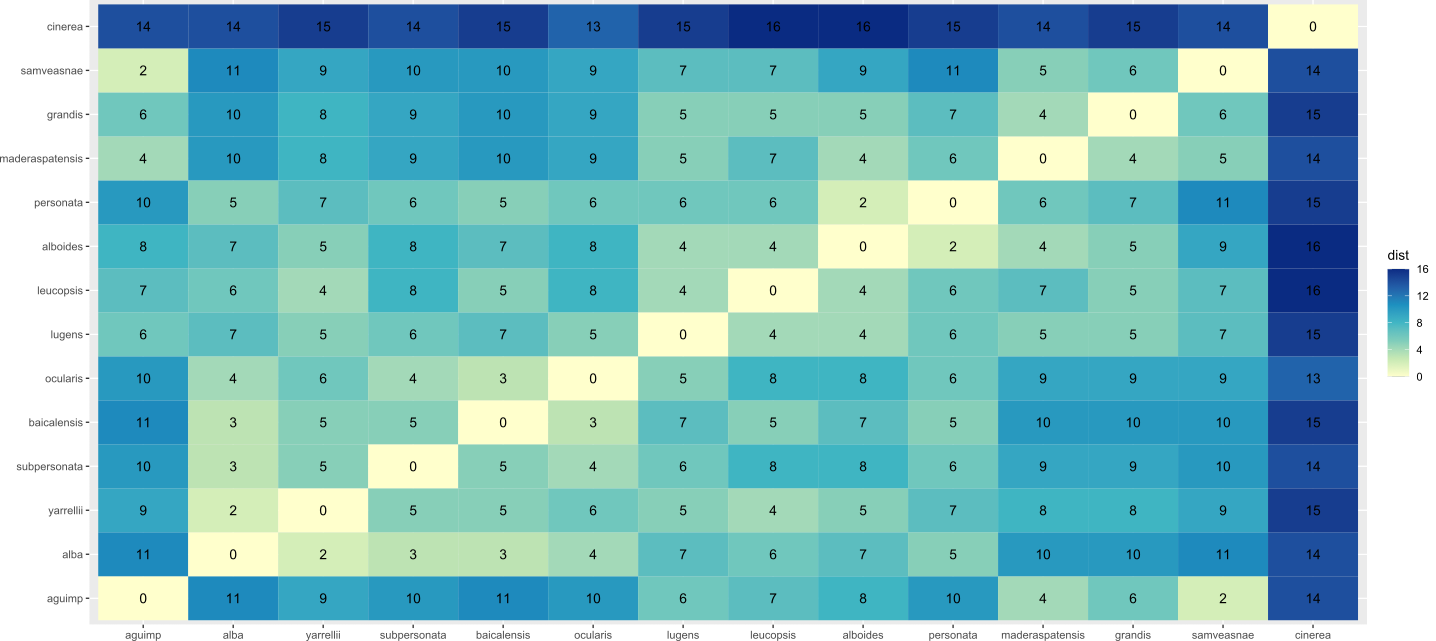

***Figure S1.*** *Distance matrix calculated based on 16 plumage characteristics scored in breeding males.*

**Table S2.** List of the samples used in this study.

| **Sample ID** | **Museum number** | **Species** | **Locality** | **Sex** |
| --- | --- | --- | --- | --- |
| Mot_agu_agu_01P12604_106_S6 | 95556 / GAV1132 | *Motacilla aguimp* | 95 km S, 45 km W of Kimberley, Northern Cape Prov., South Africa | Male |
| Mot_agu_agu_02 P12604_107_S7 | 95558 / GAV1813 | *Motacilla aguimp* | 20 km NW Kimberley, Northern Cape Prov., South Africa | Male |
| Mot_agu_vid_01 P12604_108_S8 | 110295 /JK 02 447 | *Motacilla aguimp* | Mulanje Massif Forest Reserve Lodge, Malawi | Male |
| Mot_agu_vid_02 P12604_109_S9 | 110296 /JK 02 448 | *Motacilla aguimp* | Mulanje Massif Forest Reserve Lodge, Malawi | Female |
| Mot_agu_vid_03 P12604_110_S10 | UG57 | *Motacilla aguimp* | Uganda | Female |
| Mot_agu_vid_05 P12604_112_S12 | ZMUK 132566 / TLW6-22.10.97 | *Motacilla aguimp* | Uganda | Unsexed |
| Mot_alb_alb_01 P12604_114_S14 | UWBM 61244 | *Motacilla alba* | Temryuk, Krasnodarskiy Kray, Russia | Female |
| Mot_alb_alb_02 P12604_123_S22 | UWBM 61246 | *Motacilla alba* | Temryuk, Krasnodarskiy Kray, Russia | Female |
| Mot_alb_alb_04 P12604_125_S24 | NRM 20066613 | *Motacilla alba* | Örebro, Kvismaren, Rysjön, Närke, Sweden | Male |
| Mot_alb_alb_05 P12604_126_S25 | NRM 20056584 | *Motacilla alba* | Uppsala, Sweden | Male |
| Mot_alb_alb_06 P12604_127_S26 | NRM 20056585 | *Motacilla alba* | Uppsala, Sweden | Female |
| Mot_alb_alb_07 P12604_128_S27 | NRM 20026381 | *Motacilla alba* | Gräsö, Örskär, Uppland, Sweden | Male |
| Mot_alb_alb_08 P12604_129_S28 | NRM 20006306 | *Motacilla alba* | Stockholm, Sweden | Male |
| Mot_cin_cin_01  P12604_117_S17 | UWBM 44583 / SAR 6222 | *Motacilla cinerea* | Sokhoch, Kamchatka, Russia | Male |
| Mot_cin_cin_04  P12604_120_S20 | UWBM 75848 / MSUZM 2000/N302 | *Motacilla cinerea* | Mugur-Aksy, Respublika Tyva, Russia | Female |
| Mot_cin_cin_05 P12604_121_S21 | ZMUK 142023 / MWA4-12.4.07 | *Motacilla cinerea* | Kok Meinok, Ala Too mts, Kyrgyzstan | Unsexed |
| Mot_cin_cin_06 P12604_282_S54 | ZMUK 148951 / JBK25-13.2.13 | *Motacilla cinerea* | Denmark | Unsexed |
| Mot_cin_cin_07 P12604_283_S55 | IOZ 10592 / SX 009 | *Motacilla cinerea* | Erlang village, HuangbaiyuaTown, Taibai county, Shaanxi, China | Unsexed |
| Mot_cin_cin_08 P12604_284_S56 | NRM GMN 2008-21900 / GNM AvSu11260 (DNA 24794) | *Motacilla cinerea* | Horred, Halland, Sweden | Unsexed |
| Mot_gra_nan_02 P12604_152_S50 | YIO 1995-0065 | *Motacilla grandis* | Hiroshima-city, Hiroshima Pref., Japan | Unsexed |
| Mot_gra_nan_03 P12604_153_S51 | YIO 1995-0069 | *Motacilla grandis* | Hiroshima-city, Hiroshima Pref., Japan | Unsexed |
| Mot_gra_nan_04 P12604_154_S52 | YIO 2006-5292 | *Motacilla grandis* | Tsukuba-city, Ibaraki, Japan | Unsexed |
| Mot_gra_nan_06 P12604_247_S46 | YIO 1995-0149 | *Motacilla grandis* | Hidaka-gun, Wakayama Pref., Japan | Unsexed |
| Mot_gra_nan_07 P12604_248_S47 | YIO 1995-0150 | *Motacilla grandis* | Hidaka-gun, Wakayama Pref., Japan | Unsexed |
| Mot_gra_nan_08 P12604_249_S48 | YIO 1995-0151 | *Motacilla grandis* | Hidaka-gun, Wakayama Pref., Japan | Unsexed |
| Mot_sam_nan_02 P12604_212_S17 | NRM SAM-33 / Per (010213-3) | *Motacilla samveasnae* | Stung Treng, Cambodia | Male |
| Mot_sam_nan_04 P12604_215_S20 | NRM SAM-67 / Per (010216-7) JWDKH13 | *Motacilla samveasnae* | Stung Treng, Cambodia | Female |
| Mot_mad_nan_01 AMNH778808 | AMNH 778808 | *Motacilla maderaspatensis* | Mheskatri, Gujarat, India | Male |
| Mot_mad_nan_02 BMNH1949 | BMNH 1949.25.3010 | *Motacilla maderaspatensis* | India | Unsexed |

**Table S3.** Detailed results of the D statistics.

| **P1** | **P2** | **P3** | **N_BBAA_** | **N_ABBA_** | **N_BABA_** | ***D*** | **p-value** | **Z score** |
| --- | --- | --- | --- | --- | --- | --- | --- | --- |
| *grandis* | *alba* | *aguimp* | 444507 | 479744 | 358611 | 0.14 | 0 | 44.25 |
| *samveasnae* | *alba* | *aguimp* | 421802 | 520800 | 363900 | 0.18 | 0 | 34.34 |
| *samveasnae* | *grandis* | *aguimp* | 501569 | 433148 | 396942 | 0.04 | 0 | 9.44 |
| *samveasnae* | *grandis* | *alba* | 442224 | 459395 | 395500 | 0.07 | 0 | 13.28 |

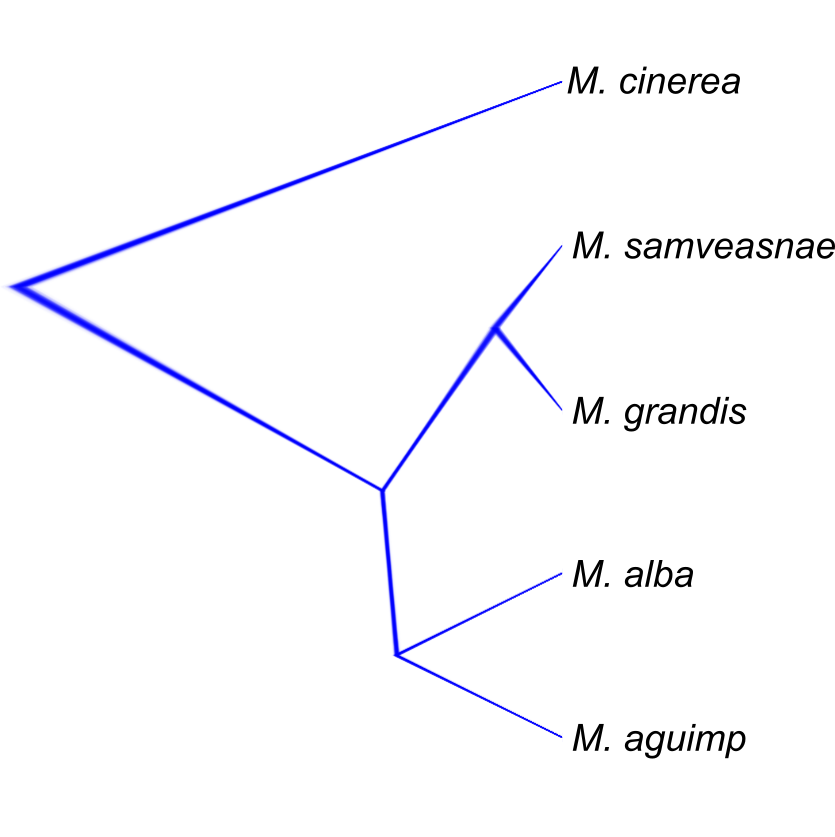

***Figure S2.*** *Snapper species tree inferred from the* snapper_4sp *dataset (182,693 SNPs).*

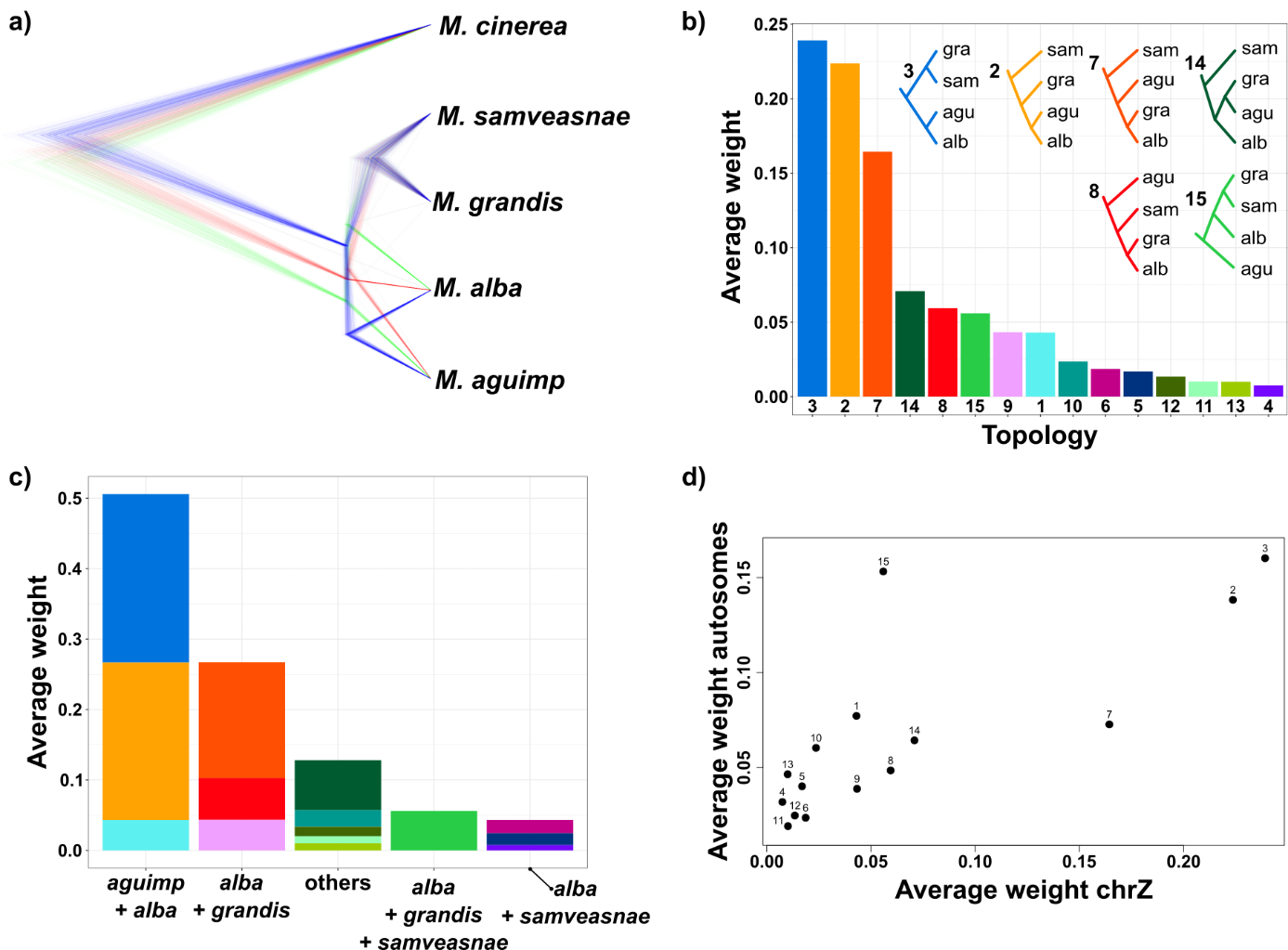

***Figure S3.*** *Phylogenetic analyses of the Z chromosome. a) snapper species tree based on 14,904 unlinked SNPs. b) Average weight of the 15 possible topologies based on 50 kb non-overlapping sliding windows. c) The same data with the topologies aggregated in five categories depending on the nodes the display (colors in b and c are the same as in Figure 4). d) Relationships between the average weights of the 15 topologies on the autosomes and Z chromosome.*

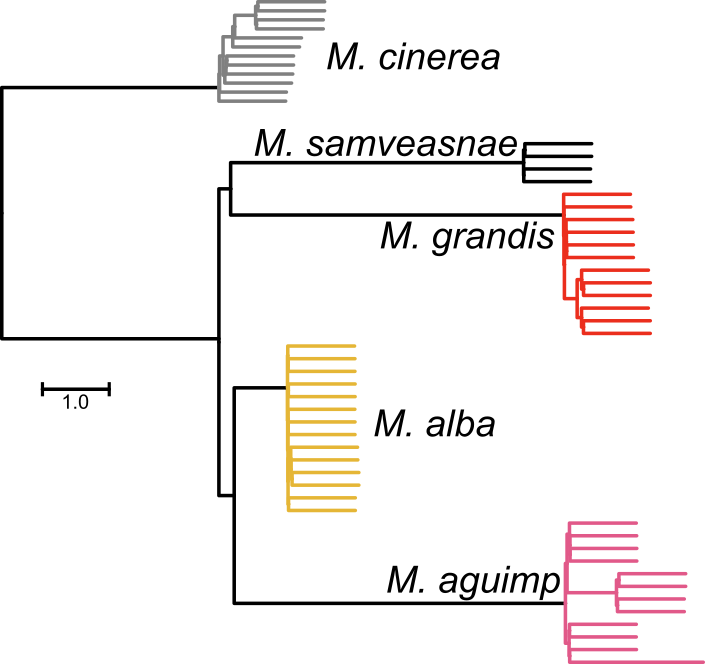

***Figure S4.*** *ASTRAL species tree inferred from the* ASTRAL_4sp *dataset (9,443 trees calculated in non-overlapping and non-adjacent 50kb sliding windows).*

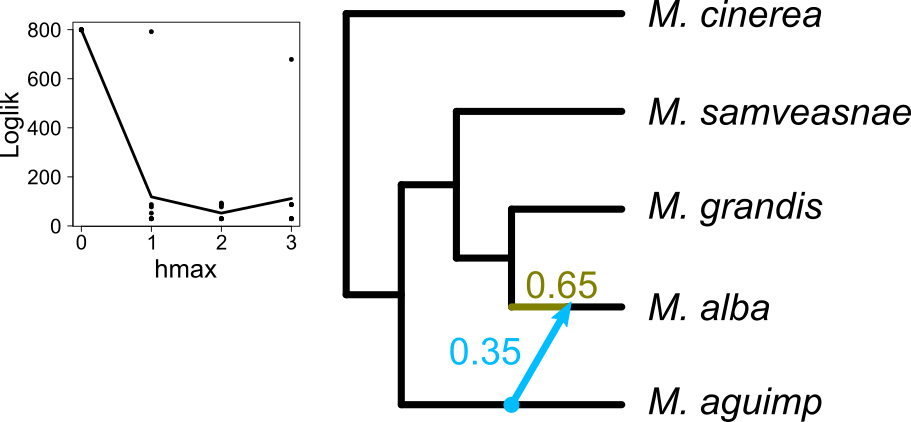

***Figure S5.*** *Phylogenetic network inferred from the* snapper_4sp *dataset (182,693 SNPs). The graph on the left shows the changes in loglik when increasing* hmax. *The green edge and blue arrow show the major and minor edges, respectively, of the inferred hybridization event, and the associated numbers are their inheritance probabilities (γ).*

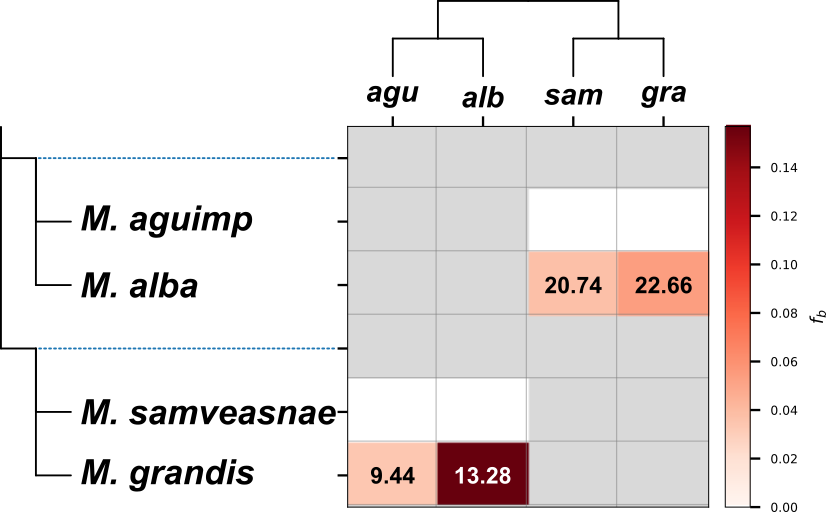
***Figure S6.*** f_b_ *statistics calculated based on an alternative topology. Numbers in the cells indicate Z scores.*

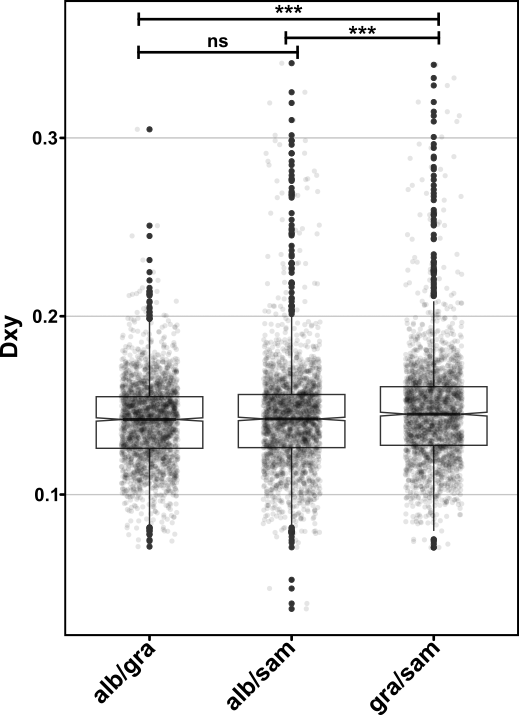

***Figure S7.*** *D*_XY_ *between* M. alba *and* M. grandis, M. alba *and* M. samveasnae, *and* *M*. *grandis and* *M*. *samveasnae in non-introgressed 50 kb sliding windows.*
